# Characterising memory loss in patients with autoimmune limbic encephalitis hippocampal lesions

**DOI:** 10.1101/579979

**Authors:** Meher Lad, Sinéad L. Mullally, Alexandra L. Houston, Tom Kelly, Timothy D. Griffiths

## Abstract

Since the publication of Scoville and Milner’s (1957) seminal paper, the precise functional role played by the hippocampus in support of human memory has been fiercely debated. For instance, the single question of whether the hippocampus plays a time-limited or an indelible role in the recollection of personal memories led to a deep and tenacious schism within the field. Similar polarisations arose between those who debated the precise nature of the role played by the hippocampus in support of semantic relative to episodic memories and in recall/recollection relative to familiarity-based recognition. At the epicentre of these divisions lies conflicting neuropsychological findings. These differences likely arise due to the consistent use of heterogeneous patient populations to adjudicate between these positions. Here we utilised traditional neuropsychological measures in a homogenous patient population with a highly discrete hippocampal lesion (i.e. patients with voltage-gated potassium channel complex antibody associated limbic encephalitis (VGKC-LE)). We observed impairment of recent but not remote episodic memory, a preservation of semantic memory, and recall but not recognition memory deficits. We conclude that this increasingly well-characterised group of patients may represent an important homogeneous population in which the functional role played by the hippocampus may be more precisely delineated.

---

The human hippocampus literature is abound with contentious debates and conflicting evidence surrounding the precise role played by the hippocampus in support of memory. Key areas of debate include the hippocampus’ role in recent versus remote memories (for recent reviews see Squire, Genzel, Wixted, & Morris, 2015; Sekeres, Winocur, & Moscovitch, 2018), in autobiographical versus semantic memories (for recent review see Dede & Smith, 2016), and in recollection-versus familiarity-based retrieval (for recent review see Aggleton & Morris, 2018).

With respect to the former of these debates, and in a comprehensive review of the literature from 1957 to 2010, Winocur, Moscovitch, and Bontempi (2010) reported roughly equal number of cases where hippocampal damage was either 1) associated with either non-graded or temporally-extensive retrograde amnesia or 2) resulted in the classic temporally-graded retrograde amnesia described by the Standard Consolidation (SC) model (e.g. Squire, 1992). Proponents of SC model have attributed the source of this variation to lesions selectivity, whereby ungraded retrograde amnesia is a consequence of extra-hippocampal damage. On the other side, those who argue in favour of ungraded-retrograde amnesia contend that traditional tasks of remote retrograde memory are not sensitive enough to detect subtle differences between the recall of true remote autobiographical memories and more schematic versions of such memories (for review see Winocur & Moscovitch, 2011). Hence, according to this perspective, traditional findings of preserved remote memories are likely to be an artefact of limited methodologies.

Relatedly, the neuropsychological evidence for and against a unified role for the hippocampus in both semantic and episodic memory is complex (Winocur et al., 2010; Winocur & Moscovitch, 2011). As SC model does not distinguish between the role played by the hippocampus in support of different forms of declarative memories (e.g. between episodic/context-dependent memories and semantic/context-general memories), neuropsychological cases and/or group studies that appear to demonstrate a dissociation between these different forms of memories have attracted much theoretical discussion. For instance, the seemingly preserved semantic memory coupled with the severely impaired episodic memory reported in cases of selective hypoxic damage to the hippocampus in childhood (e.g. Vargha-Khadem et al., 1997) are at odds with the form of hippocampal-equivalence proposed by the SC model. Moreover, reports of spared semantic memory within a context of severely impaired episodic memory in patients with adult-onset MTL/HC amnesia (e.g. Rosenbaum et al., 2008; Verfaellie, Koseff, & Alexander, 2000) are difficult to accommodate within such models. However, the structural selectivity of these dissociations have been fiercely disputed and counter-findings are frequently presented (e.g. Manns, Hopkins, & Squire, 2003; Reed & Squire, 1998; but see Winocur & Moscovitch, 2011). Moreover, uncertainties surrounding a clear division between episodic and semantic memory are evident in the semantic dementia literature (e.g. Greenberg & Verfaellie, 2010; Burianova, McIntosh, & Grady, 2010).

Further divisions within the field also exist between so-called dual-process memory theorists (for reviews see Aggleton & Brown, 2006; Eichenbaum, Yonelinas, & Ranganath, 2007; Gardiner & Java, 1993; Montaldi & Mayes, 2010; Yonelinas, 2002) and single-process memory theorists (for reviews see Clark, 2018; Squire, Wixted, & Clark, 2007; Wixted, 2007). In essence, dual-process theories argue that the extended hippocampus (i.e. the hippocampus, the anterior nucleus of the thalamus, and the mamillothalamic tract) is selectively involved in recollection-based memory processes, whilst familiarity-based recognition is supported by the perirhinal memory system (i.e. the perirhinal cortex and mediodorsal nucleus). Recollection-based memory is defined as memory with an associated subjective feeling of remembering, which encompasses both free recall and event recognition if that recognition is accompanied by the full recall of the event and encoding context. Familiarity-based recognition is recognition that occurs with an isolated sense of familiarity and in the absence of full recollection. Findings consistent with this viewpoint offer evidence of specific recollection deficits in patients with selective damage to the hippocampal memory system (e.g. Brandt, Gardiner, Vargha-Khadem, Baddeley, & Mishkin, 2009; Holdstock et al., 2002) and/or a double dissociation between recall- and familiarity-based recognition (e.g. Bowles et al., 2010; see also Brandt, Eysenck, Nielsen, & von Oertzen, 2016). However, single-process theorists dispute these functional and structural dissociations, arguing that these apparent dissociations are driven by differences in memory strength, and point to the studies that report impairment of both recall and recognition following bilateral hippocampal damage (e.g. Manns et al., 2003).

The extent of the conflict in the above studies is striking, not least due to the fact that many have been observed using standardised neuropsychological tasks such as the Autobiographical Memory Interview (AMI) (Kopelman, Wilson, & Baddeley, 1990) and the Doors and People Test (D&P) (Baddeley & Nimmo-Smith, 1994). Standardised memory tasks, if administered correctly, eliminate variation in administration and scoring methodology as a source of these differences. Moreover, the D&P test equates the difficulty levels of the recall and recognition subtests - a control that counters the memory strength hypothesis proposed by single theorists to explain apparent recall/recognition dissociations in the literature (e.g. Dunn, 2008). Hence, reconciling these theoretical stalemates using identical neuropsychological methods has been largely unsuccessful. Moreover, whilst novel neuroimaging techniques such as the combination of high-resolution structural and functional magnetic resonance imaging and advanced analytical methods (such as multi-voxel pattern analysis) can undoubtedly add unique leverage on these issues (for review see Maguire, 2014), traditional neuropsychological approaches still play an important role in the resolution of these conflicts.

One persistent and major challenge for those undertaking such studies is that patients with selective and uniform hippocampal damage are exceedingly rare. Hence, group studies with hippocampal amnesic patients typically utilise patients with damage acquired through a range of diverse aetiologies (e.g. cardiac arrest, carbon monoxide poisoning, drug overdose, or ‘unknown’, Manns et al., 2003) or include patients with varying neuropathologies including damage to the hippocampus, frontal lobe and thalamus (e.g. Manns & Squire, 2002). These studies therefore assume a uniformity across patients that is largely unsubstantiated, as studies combining both neuropsychological observations and post-mortem descriptions are understandably rare (for notable exceptions see Zola-Morgan, Squire, & Amaral, 1986; Remple-Clower, Zola, Squire, & Amaral, 1996; Annese et al., 2014), and determining hippocampal functionality on the basis of structural MRI can be misleading (Mullally, Hassabis & Maguire, 2012).

In this study, we assess a homogenous population with a highly discrete hippocampal lesion that is clinically stable after an acute phase: patients with voltage-gated potassium channel complex antibody associated limbic encephalitis (VGKC-LE) (Miller et al., 2017; Finke et al, 2016). VGKC-LE is a rare autoimmune condition discovered in 2004 with a prevalence of about 1 in 400,000 (Vincent et al., 2004). It is an autoimmune inflammatory disorder that causes long-term memory impairment, seizures and sometimes-behavioural disturbances in its acute phase, but patients recover and can be left with selective memory deficits (Buckley et al., 2001). Patients who test positive for VGKC-complex antibodies are further subdivided into those with anti leucine-rich glioma inactivated (anti-LGI-1) encephalitis (who present with limbic symptoms), anti contactin-associated protein-like 2 (anti CASPR-2) (who present with both central and peripheral symptoms) and a third group who do not have antibodies against LGI-1 or CASPR-2 and who present with heterogeneous symptoms (Bastiaansen, van Sonderen, & Titulaer, 2017). Both LGI-1 and CASPR-2 are different, well-described clinical phenotypes (van Sonderen, Petit-Pedrol, Dalmau & Titulaer, 2017), with the former specific for the hippocampus (van Sonderen et al., 2016). Here we consider patients with this brain phenotype as a unique opportunity to test the effect of an anatomically-selective hippocampal lesion. Hence, unlike the previously discussed neuropsychological studies of patients with selective hippocampal damage, we utilised a uniform cohort of stable patients to adjudicate between the entrenched and conflicting theoretical perspectives described above. In total, we tested seven patients (two female, mean age: 66 years, range: 51-70 years) with VGKC-LE recruited via the Cognitive Clinic at the Royal Victoria Infirmary, Newcastle upon Tyne, United Kingdom. Patients were selected if they had positive VGKC-complex antibody level of >1000pM at the time of diagnosis and a clinical phenotype consistent with LGI-1 limbic encephalitis after review by a Cognitive Neurologist (see Table S1 for further details).

At presentation in the acute phase of illness, MR brain imaging showed increased hippocampal signal intensity on T2 or FLAIR sequences in 5 out of 7 patients (Figure. 1A: Acute Phase). Subsequently, in the stable chronic phase and more than 1 year after acute presentation, all patients had additional structural MRI (see supplemental Methods). This revealed hippocampal atrophy in each patient that was specific to hippocampal, as opposed to parahippocampal structures (Figure 1A: Stable Phase). An extensive neuropsychological assessment was also performed within 2 months of patients undergoing stable phase structural MRI. Results were consistent with a selective memory impairment as opposed to a global cognitive insult.

**Figure 1.**
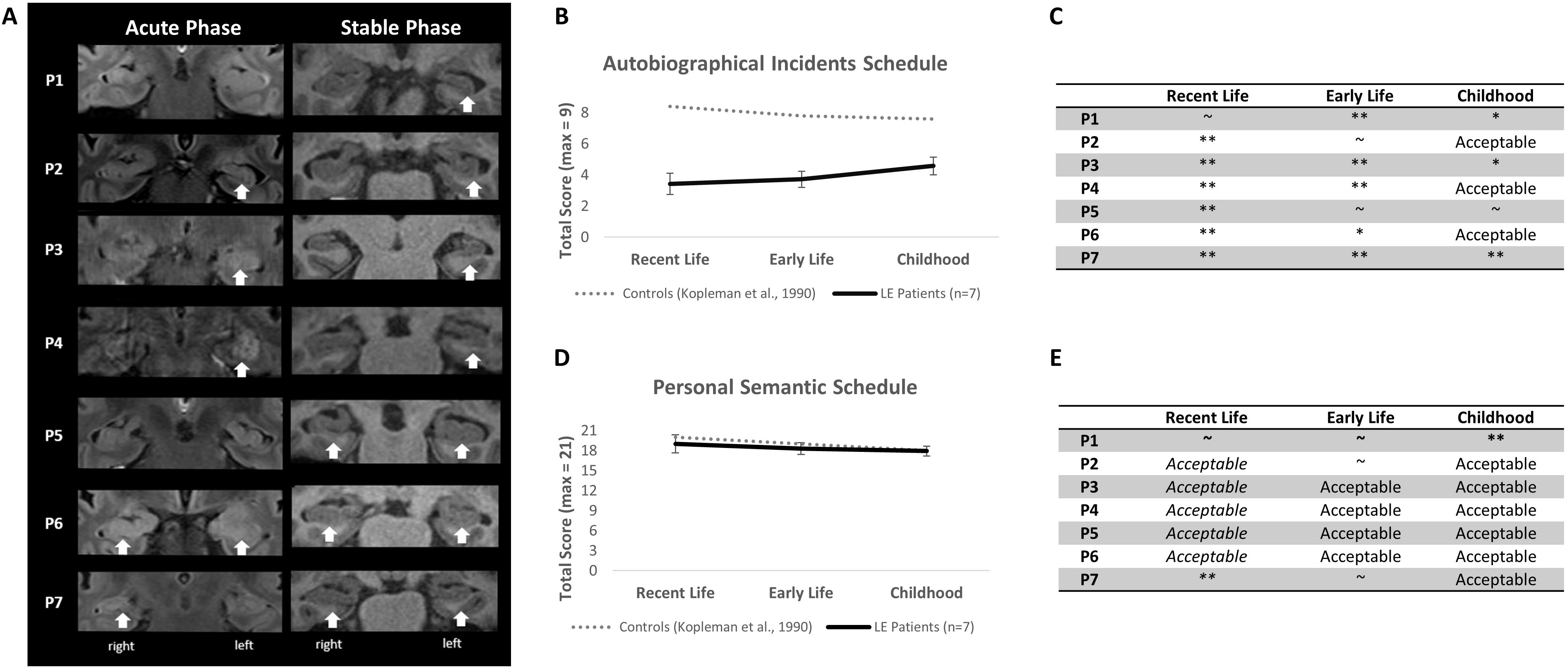
**A)** Structure MRI of patients from initial presentation in the acute phase (T2 coronal FLAIR; left column) and at time of neuropsychological testing in the acute phase (T1 coronal; right column). White arrows indicate increased signal intensity in the left column and subsequent atrophy in the right column (as reported by neuroradiologists). **B)** The Autobiographical Memory Interview – Autobiographical Incidents (group data). VGKC-LE patient scores are represented relative to the cut-off points for healthy controls cited in Kopleman et al. (1990). NS = non-significant differences between epochs. **C**) The AMI: Autobiographical Incidents (individual patient data). ** Definitely Impaired (scores at or below which none of the healthy controls scored); * Probably Impaired (> 2 StDev below the control mean); ∼ Borderline (between 1 StDev – 2 StDev below the control mean); Acceptable (±1 StDev of the control mean). **D**) AMI: Personal Semantic Schedule (group data). VGKC-LE patient scores are represented relative to the cut-off points for healthy controls cited in Kopleman et al. (1990). NS = non-significant differences between epochs. **C)** The AMI: Personal Semantic Schedule (individual patient data). ** Definitely Impaired (scores at or below which none of the healthy controls scored); * Probably Impaired (> 2 StDev below the control mean); ∼ Borderline (between 1 StDev – 2 StDev below the control mean); Acceptable (±1 StDev of the control mean).

To test the outstanding issues outlined above, we administered two further standardised neuropsychological measures - the AMI and the D&P test (Baddeley, 1994). The AMI is a standardised test of autobiographical and personal semantic memories that uses a structured interview to assess memory across three different time-periods (i.e. recent events: within the last year; early adulthood: 19-29 years old; and childhood: up until 18 years of age). The D&P assesses verbal and visual recall and recognition and critically, difficulty levels of the recall and recognition subsets is equated. Both the AMI and the D&P tasks have been used extensively in the hippocampal literature enabling direct comparison with previous work.

With respect to the AMI, we hypothesised that if the hippocampus plays a time-limited role in memory retrieval, then VGKC-LE patients should show impairment on recall of recent autobiographical memories but a preservation of remote memories. However, if retrograde memory deficits are evident across the lifespan (and are therefore non-graded), then this results would favour alternative theories, such as Multiple Trace Theory (MTT; Nadel & Moscovitch, 1997), Transformation Theory (Winocur & Moscovitch, 2011) and Scene Construction Theory (Maguire & Mullally, 2013), that do not draw this distinction. Similarly, the AMI also enabled us to assess whether VGKC-LE patients would be equally impaired on autobiographical and semantic memory (as predicted by the standard consolidation model), or show greater impairment to autobiographical memories (as predicted by the above alternative theories).

As anticipated, the VGKC-LE patients demonstrated a striking amnesia for autobiographical material, with six of the seven patients showing definite impairment in the recall of recent autobiographical memories (Figure 1C). Moreover, their performance at a group level fell well below the range observed in health controls (Kopelman et al., 1990; Figure 1B). However, this clear impairment was coupled with a robust preservation of personal semantic memories at each of these time points (Figure 1D). This dissociation is consistent with the recently reported dissociation between episodic and semantic memory impairment/preservation observed in a group of 16 LGI1-VGKC-LE patients (Miller et al., 2017). Moreover, at the group level, and contrary to the standard consolidation model, our VGKC-LE patients demonstrated no temporal-gradient when recalling past personal memories. More specifically, no significant within-group differences were observed between recent and early life autobiographical memories (t(6)=-0.367, p=0.726), between early life and childhood autobiographical memories (t(6)=-1.131, p=0.301), and between recent life and childhood autobiographical memories (t(6)=-1.686, p=0.143) (Figure 1B). The same absence of a temporal-gradient was evident semantic memories (t(6)=0.977, p=0.366; t(6)=0.803, p=0.452; t(6)=0.314, p=0.764; Figure 1D).

At an individual level, the pattern of impairment of AM across each life epoch was more mixed; with three of the seven patients reporting autobiographical memories for their childhood that fell within an acceptable range (Figure 1C). Each of these cases however, resides on the lower boundary of acceptable category. The underlying reason for these qualitative (but non-significant quantitative) differences is unclear. One possibility (in line with MTT and Transformation theory) is that these acceptable childhood autobiographical memories may be disproportionally benefitting from well-rehearsed personal semantic childhood knowledge which are clearly intact in this group (see Figure 1E). Without the benefit of more nuanced measures, this modest but non-significant benefit for childhood autobiographical memories remains unclear. These findings raise important challenges for standard consolidation model.

With respect to the D&P, we asked whether VGKC-LE patients would demonstrate a uniform impairment across the recall and recognition subtests (consistent with a single-process theories) or whether they would display a dissociation (i.e. consistent with a dual-process theories). Relative to the published performance norms (Baddeley, 1994), six of the seven patients (P2-P7) scored lower on recall than recognition tasks (see Table S3). In three of these patients (P3, P6 and P7), recall was markedly-impaired and this contrasted with clearly preserved recognition scores. The final patient performed both subtests to a high level of accuracy (P01; see Table S3). To further explore the issue of task difficulty we recruited fourteen age and gender-matched controls (corresponding to two per patient; four female, mean age: 65 years, range: 52-73). Consistent with the above report, LE patients performed significantly worse than their matched controls on immediate verbal recall (*U*=2.0, *P*<0.001), delayed verbal recall (*U*=20.0, *P*=0.031), and delayed visual recall (*U*=22.0, *P*=0.046), but not on immediate visual recall (*U*=36.5, *P*=0.360). In contrast, no significant deficits in either verbal recognition (*df*= *19, t*=-0.288, *P*=0.777) and visual recognition (*df*= *19, t*=0.645, *P*=0.527) memory were observed in the patient group relative to the matched controls. Moreover, there were no differences when the easy (i.e. Set A) and hard (i.e. Set B) trials were compared between patients and controls [Verbal Recognition Set A (*U*=20.5, *p*=0.161), Set B (*df*= *15, t*=0.539, *p*=0.598), Visual Recognition Set A (*U*=26.0, *p*=0.417) and Set B (*df*= *15, t*=0.501, *p*=0.623) (Figure 2B).

**Figure 2.**
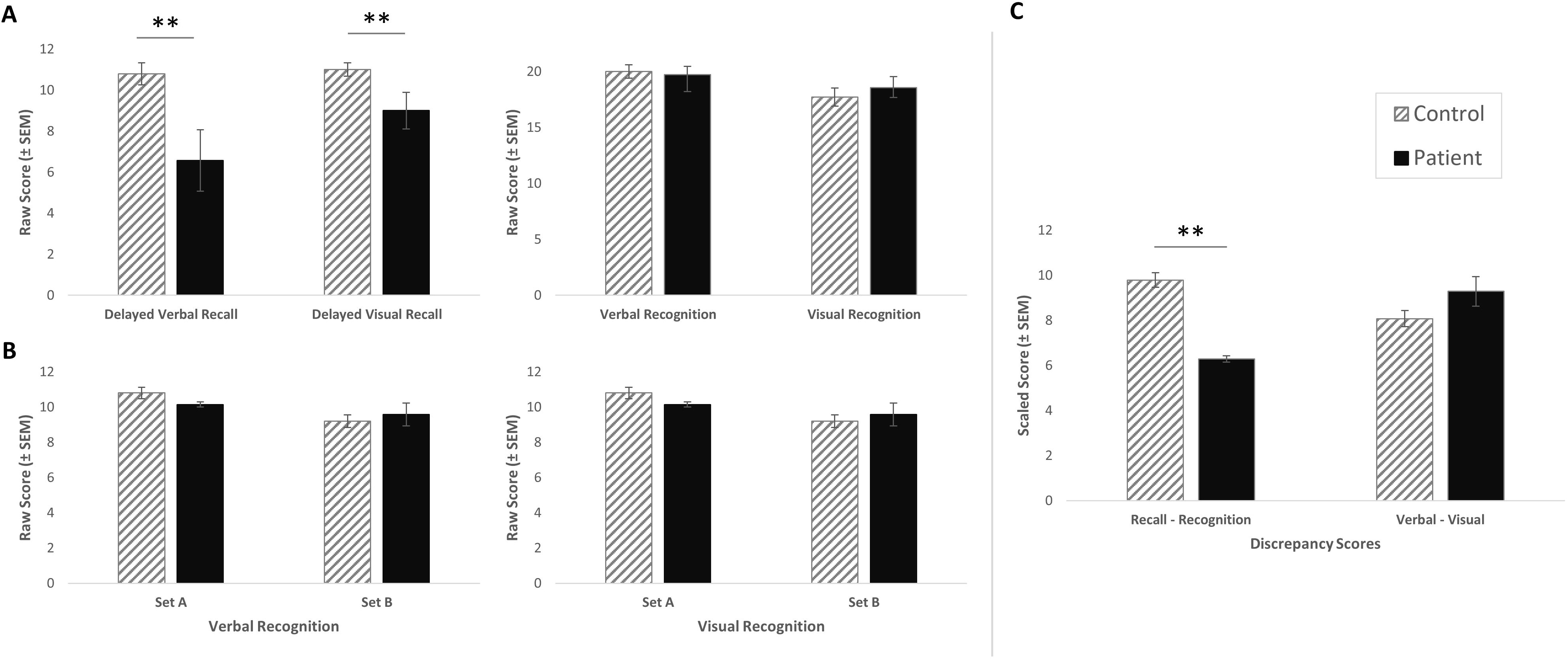
Bar graphs representing patient and matched-control raw scores in **A)** Overall Verbal versus Visual Recall Memory and Verbal versus Visual Recognition Memory. Recall is significantly lower than Recognition Memory in both domains for patients. **B)** Verbal Recognition Memory – Set A (easy) and Set B (hard) and Visual Recognition Memory – Set A (easy) and Set B (hard). **C)** Overall Verbal versus Visual Memory Measures and Recall versus Recognition Memory. Recall is significantly lower than Recognition Memory in patients (U=14.0, P=0.007). No significant difference for verbal/visual discrepancy (df= 15, t=1.580, P=0.131) was observed.

Hence, in this study we explored three key area of controversy in the memory literature, i.e. whether damage to the human hippocampus selectively impairs recent versus remote memories, autobiographical versus semantic memories, and recollection-versus familiarity-based retrieval, in stable patient population that provide a novel lesion model to test different hypotheses for hippocampal function. Further work is now necessary to explore additional hippocampal-based processes with this patient group and more studies are needed to see if *in-vivo* imaging findings correlate with discrete cognitive markers of impaired hippocampal function. However, this increasingly well-characterised group of patients may represent an important neuropsychological model of selective hippocampal impairment (perhaps even at the level of hippocampal subfields; Miller et al., 2017), which, in combination with of high-resolution structural and functional MRI and advanced analytical methods, could offer important traction on many of the entrenched impasses firmly rooted in the hippocampal literature.

## References

Aggleton, J. P., & Brown, M. W. (2006). Interleaving brain systems for episodic and recognition memory. Trends in cognitive sciences, 10(10), 455–463.

Aggleton, J. P., & Morris, R. G. (2018). Memory: Looking back and looking forward. Brain and Neuroscience Advances, 2, 2398212818794830.

Annese, J., Schenker-Ahmed, N. M., Bartsch, H., Maechler, P., Sheh, C., Thomas, N.,…& Klaming, R. (2014). Postmortem examination of patient HM’s brain based on histological sectioning and digital 3D reconstruction. Nature communications, 5, 3122.

Baddeley, A. D. E. H.; Nimmo-Smith, I. (1994). Doors and People: a test of visual and verbal recall and recognition. In. Bury St. Edmunds, England: Thames Valley Co.

Bastiaansen, A. E., van Sonderen, A., & Titulaer, M. J. (2017). Autoimmune encephalitis with anti-leucine-rich glioma-inactivated 1 or anti-contactin-associated protein-like 2 antibodies (formerly called voltage-gated potassium channel-complex antibodies). Current opinion in neurology, 30(3), 302–309.

Bowles, B., Crupi, C., Pigott, S., Parrent, A., Wiebe, S., Janzen, L., & Köhler, S. (2010). Double dissociation of selective recollection and familiarity impairments following two different surgical treatments for temporal-lobe epilepsy. Neuropsychologia, 48(9), 2640–2647.

Brandt, K. R., Gardiner, J. M., Vargha-Khadem, F., Baddeley, A. D., & Mishkin, M. (2009). Impairment of recollection but not familiarity in a case of developmental amnesia. Neurocase, 15(1), 60–65.

Brandt, K. R., Eysenck, M. W., Nielsen, M. K., & von Oertzen, T. J. (2016). Selective lesion to the entorhinal cortex leads to an impairment in familiarity but not recollection. Brain and cognition, 104, 82–92.

Buckley, C., Oger, J., Clover, L., Tüzün, E., Carpenter, K., Jackson, M., & Vincent, A. (2001). Potassium channel antibodies in two patients with reversible limbic encephalitis. Annals of Neurology: Official Journal of the American Neurological Association and the Child Neurology Society, 50(1), 73– 78.

Burianova, H., McIntosh, A. R., & Grady, C. L. (2010). A common functional brain network for autobiographical, episodic, and semantic memory retrieval. Neuroimage, 49(1), 865–874.

Clark, R. E. (2018). Current Topics Regarding the Function of the Medial Temporal Lobe Memory System

Dede, A. J., & Smith, C. N. (2016). The functional and structural neuroanatomy of systems consolidation for autobiographical and semantic memory.

Dunn, J. C. (2008). The dimensionality of the remember-know task: a state-trace analysis. Psychological review, 115(2), 426.

Eichenbaum, H., Yonelinas, A. P., & Ranganath, C. (2007). The medial temporal lobe and recognition memory. Annu. Rev. Neurosci., 30, 123–152.

Finke, C., Prüss, H., Heine, J., Reuter, S., Kopp, U. A., Wegner, F.,…& Deuschl, G. (2017). Evaluation of cognitive deficits and structural hippocampal damage in encephalitis with leucine-rich, glioma-inactivated 1 antibodies. JAMA neurology, 74(1), 50–59.

Gardiner, J. M., & Java, R. I. (1993). Recognition memory and awareness: An experiential approach. European Journal of Cognitive Psychology, 5(3), 337–346.

Greenberg, D. L., & Verfaellie, M. (2010). Interdependence of episodic and semantic memory: evidence from neuropsychology. Journal of the International Neuropsychological society, 16(5), 748– 753.

Holdstock, J. S., Mayes, A. R., Roberts, N., Cezayirli, E., Isaac, C. L., O’Reilly, R. C., & Norman, K. A. (2002). Under what conditions is recognition spared relative to recall after selective hippocampal damage in humans?. Hippocampus, 12(3), 341–351.

Kopelman, M. D., Baddeley, A. D., & Wilson, B. A. (1990). AMI: The Autobiographical Memory Interview: Manual. Harcourt Assessment.

Maguire, E. A. (2014). Memory consolidation in humans: new evidence and opportunities. Experimental physiology, 99(3), 471–486.

Manns, J. R., Hopkins, R. O., & Squire, L. R. (2003). Semantic memory and the human hippocampus. Neuron, 38(1), 127–133.

Manns, J.R., Squire, L.R. The medial temporal lobe and memory for facts and events. In: A. Baddeley, B. Wilson, and M. Kopelman (Eds.), Handbook of Memory Disorders, 2nd Edition. New York: John Wiley & Sons, Ltd., 2002, 81–99.

Miller, T. D., Chong, T. T., Aimola Davies, A. M., Ng, T. W., Johnson, M. R., Irani, S. R., Vincent, A., Husain, M., Jacob, S., Maddison, P., Kennard, C., Gowland, P. A., & Rosenthal, C. R. (2017). Focal CA3 hippocampal subfield atrophy following LGI1 VGKC-complex antibody limbic encephalitis. Brain.

Montaldi, D., & Mayes, A. R. (2010). The role of recollection and familiarity in the functional differentiation of the medial temporal lobes. Hippocampus, 20(11), 1291–1314.

Maguire, E. A., & Mullally, S. L. (2013). The hippocampus: a manifesto for change. Journal of Experimental Psychology: General, 142(4), 1180.

Mullally, S. L., Hassabis, D., & Maguire, E. A. (2012). Scene construction in amnesia: An fMRI study. Journal of Neuroscience, 32(16), 5646–5653.

Nadel, L., & Moscovitch, M. (1997). Memory consolidation, retrograde amnesia and the hippocampal complex. Current opinion in neurobiology, 7(2), 217–227.

Reed, J. M., & Squire, L. R. (1998). Retrograde amnesia for facts and events: findings from four new cases. Journal of Neuroscience, 18(10), 3943–3954.

Rempel-Clower, N., Zola, S.M., Squire, L.R., and Amaral, D.G. (1996). Three cases of enduring memory impairment following bilateral damage limited to the hippocampal formation. J. Neurosci. 16, 5233–5255.

Rosenbaum, R. S., Moscovitch, M., Foster, J. K., Schnyer, D. M., Gao, F., Kovacevic, N.,…& Levine, B. (2008). Patterns of autobiographical memory loss in medial-temporal lobe amnesic patients. Journal of Cognitive Neuroscience, 20(8), 1490–1506.

Scoville, W. B., & Milner, B. (1957). Loss of recent memory after bilateral hippocampal lesions. Journal of neurology, neurosurgery, and psychiatry, 20(1), 11.

Sekeres, M. J., Winocur, G., & Moscovitch, M. (2018). The hippocampus and related neocortical structures in memory transformation. Neuroscience letters.

Squire, L. R. (1992). Memory and the hippocampus: a synthesis from findings with rats, monkeys, and humans. Psychological review, 99(2), 195.

Squire, L. R., Genzel, L., Wixted, J. T., & Morris, R. G. (2015). Memory consolidation. Cold Spring Harbor perspectives in biology, 7(8), a021766.

Squire, L. R., Wixted, J. T., & Clark, R. E. (2007). Recognition memory and the medial temporal lobe: a new perspective. Nature Reviews Neuroscience, 8(11), 872.

Vargha-Khadem, F., Gadian, D. G., Watkins, K. E., Connelly, A., Van Paesschen, W., & Mishkin, M. (1997). Differential effects of early hippocampal pathology on episodic and semantic memory. Science, 277(5324), 376–380.

Verfaellie, M., Koseff, P., & Alexander, M. P. (2000). Acquisition of novel semantic information in amnesia: effects of lesion location. Neuropsychologia, 38(4), 484–492.

Vincent, A., Buckley, C., Schott, J. M., Baker, I., Dewar, B. K., Detert, N.,…& Lang, B. (2004). Potassium channel antibody-associated encephalopathy: a potentially immunotherapy-responsive form of limbic encephalitis. Brain, 127(3), 701–712.

van Sonderen, A., Thijs, R. D., Coenders, E. C., Jiskoot, L. C., Sanchez, E., De Bruijn, M. A.,…& Titulaer, M.J. (2016). Anti-LGI1 encephalitis Clinical syndrome and long-term follow-up. Neurology, 10–1212.

Van Sonderen, A., Petit-Pedrol, M., Dalmau, J., & Titulaer, M. J. (2017). The value of LGI1, Caspr2 and voltage-gated potassium channel antibodies in encephalitis. Nature Reviews Neurology, 13(5), 290.

Winocur, G., Moscovitch, M., & Bontempi, B. (2010). Memory formation and long-term retention in humans and animals: Convergence towards a transformation account of hippocampal–neocortical interactions. Neuropsychologia, 48(8), 2339–2356.

Winocur, G., & Moscovitch, M. (2011). Memory transformation and systems consolidation. Journal of the International Neuropsychological Society, 17(5), 766–780.

Wixted, J. T. (2007). Dual-process theory and signal-detection theory of recognition memory. Psychological review, 114(1), 152.

Yonelinas, A. P. (2002). The nature of recollection and familiarity: A review of 30 years of research. Journal of memory and language, 46(3), 441–517.

Zola-Morgan, S., Squire, L.R., and Amaral, D.G. (1986). Human amnesia and the medial temporal region: Enduring memory impairment following a bilateral lesion limited to field CA1 of the hippocampus. J. Neurosci. 6, 2950–2967.

