## Supplemental Information for "Characterising memory loss in patients with autoimmune limbic encephalitis hippocampal lesions"

*Patient Background and Treatment*

Patients with VGKC-LE were recruited via the Cognitive Clinic in Newcastle upon Tyne, United Kingdom. Patients’ records were reviewed, between May and December 2015, for positive serum samples for VGKC antibodies and the classical clinical syndrome for LGI-1 VGKC-LE (see Table S1). All patients selected had a subacute deterioration over a period of weeks to months with retrograde and anterograde amnesia, with or without seizures, as the presenting complaint. Routine blood tests excluded infectious and metabolic causes of memory loss. All patients had a lumbar puncture to exclude infectious encephalitis and had an acellular tap with protein and glucose within acceptable parameters. Patients received inpatient treatment with antiepileptic drugs and immunomodulatory therapies that included steroids, intravenous immunoglobulins (IVIG), plasma exchange and long-term immunosuppression. Patients with significant co-morbidity, those requiring active management, those with significant small vessel disease burden on T2 weighted MRI and those with an atypical phenotype were excluded from this study. Fifteen patients were initially identified and seven met the above criteria for participation.

**Table S1**

| ***Table S1* Clinical details of patients recruited to current study** | | | | | |
| --- | --- | --- | --- | --- | --- |
| **Patient Number** | **Age, Sex** | **Clinical Features** | **VGKC Antibody Level** | **Treatment** | **Time after tested** |
| **1** | 70M | Am, Sz | 2469 | PEX, St, AED | 5 years |
| **2** | 51M | Am, Sz | 1604 | PEX, St, AED | 1 year |
| **3** | 73F | Am, FBD | 4845 | PEX, St, AED | 3 years |
| **4** | 75F | Am, FBD | 1745 | IVIG, St, Imm | 7 years |
| **5** | 62M | Am, Sz | 1065 | St, AED | 4 years |
| **6** | 63M | Am, Sz | 2663 | St, Imm, AED | 6 years |
| **7** | 69M | Am, Sz | 1001 | St, AED | 5 years |
| Am – Amnesia, Sz – Seizures, FBD – Faciobrachial dystonic seizures, PEX – Plasma Exchange, St – Steroids, AED – Antiepileptic Drug, Imm – Immunosuppression, IVIG – Intravenous Immunoglobulin, M – Male, F – Female | | | | | |

*Additional structural MRI on Patients*

Additional sMRI was performed using a 1.5T Philips NT Intera scanner with a 5-channel phased array head coil. High resolution anatomical scans (3D T1 gradient echo, 180 slices, 0.94 mm × 0.94 mm × 0.94 mm plane resolution and slice thickness, TR = 14ms, TE = 6.5ms, flip angle = 22°) were acquired for each patient.

*Additional Clinical Neuropsychological Assessments on Patients*

*Neuropsychological Characterisation*

As part of their routine neuropsychological evaluation patients performed cognitive tests to assess general intelligence (IQ), executive function, visuospatial ability, and memory (retrograde and anterograde memory). Current wellbeing was also assessed.

*General Intelligence*. An estimate of pre-morbid IQ was generated using the Weschler Test of Adult Reading (WTAR) (Wechsler, 2001). An index-based, seven subtest, short form of the Weschler Adult Intelligence Scale III (WAIS-III) was administered to estimate current IQ (Crawford *et al.*, 2008). The seven subtests were as follows: Vocabulary, Similarities, Block Design, Matrix Reasoning, Arithmetic, Digit Span, and Digit Symbol - Coding. Scores were computed from an executive program which produced index scores, confidence intervals and the reliability and abnormality of the differences between index scores.

*Executive Function*: Executive function was assessed using the Delis-Kaplan Executive Function System (DKEFS): Verbal Fluency Test, Colour-Word Interference Test, and the Trail Making Test (Delis, et al., 2004), and the Hayling and Brixton tests (Burgess & Shallice, 1997; (Bielak, et al., 2006) the Hayling Sentence Completion Test and the Brixton Spatial Anticipation Test.. A DKEFS index was calculated as a composite score of executive function (Crawford, Garthwaite, Sutherland, & Borland, 2011). The Hayling-Brixton scores were converted to their IQ equivalents according to the manual.

*Visuospatial Ability:* Visuospatial ability was assessed using two tests from the Visual Object and Space Perception test (VOSP); the Cube Analysis test and the Object Decision test (E. K. J. Warrington, M., 1991).

*Anterograde memory*: Anterograde memory was assessed using the British-normed BIRT Memory and Information Processing Battery (BMIPB; Coughlan, Oddy and Crawford, 2007; Story Recall (immediate and delayed), Figure Recall (copy, immediate and delayed recall) and List Learning, and the Warrington Recognition Memory Test (E. K. Warrington, 1984); Words and Faces). The BMIPB was administered as there are British norms available. These tasks share a number of similarities with more commonly used measures of memory; i.e. Story Recall is analogous with the Wechsler Memory Scale Logical Memory subtest, Figure Recall is analogous to the Rey Complex Figure Test (Rey, 1941), and List Learning (List A x5 (max); List B; List A) is roughly analogous to the Rey Auditory and Verbal Learning Task (Cohen, 1996).

**Additional Methods: The Doors and People Test**

In a separate testing session, the Doors and People (Baddeley, 1994) test of recall and recognition memory was administered to all patients. The test consists of sections testing verbal and visual memory in the domains of recall and recognition memory, each designed to be of equal difficulty based on a large group of healthy participants. For *verbal recall* (People Test), four photographs showing an individual with their printed name and occupation are presented on separate cards. After the fourth picture is viewed, a participant is asked to recall each name cued by their profession. This is repeated until all four names are recalled correctly or for three trials. For *visual recall* (Shapes Test), participants copy four line drawings one at a time. They are then asked to re-draw the four shapes from memory. Again, the procedure was repeated until they correctly recalled all items or a maximum of three trials. The *verbal recognition* (Names Test) consists of two subsets A (easy) and B (hard). In set A, 12 female first and surnames are presented on cards for three seconds each and read aloud. Immediately afterwards, participants are asked to select a name from a list of four distractor names. Set B is then administered and consists of male names but is harder in that the foils differ in only one syllable in the surname. In the *visual recognition* (Doors Test) task, there are two subsets A and B consisting of photographs of 12 doors. Each is presented sequentially for three seconds. Thereafter, a participant views arrays of four doors and chooses one. Set B is harder as the doors appear to be more similar in each array.

The Doors and People test allows for additional metrics to be calculated after converting raw scores to scaled measures. These are overall verbal and visual scores, verbal/visual discrepancy score (difference between verbal recall and recognition scores and visual recall and recognition scaled scores), overall recall and recognition scores and recall/recognition discrepancy (difference between total recall and total recognition scaled scores) scores.

In addition, fourteen age and gender-matched controls (corresponding to two per patient; four female, mean age: 65 years, range: 52-73) completed the Doors and People Task. Control participants were recruited via the participant database at the Institute of Neuroscience, Newcastle University, Newcastle upon Tyne, United Kingdom. They were closely matched to the patients for age (*df*= *19, t*=0.236, *P*=0.816), sex and Weschler Test of Adult Reading (WTAR) scores (Wechsler, 2001) (*df*= *19, t*=-2.04, *P*=0.056). Control participants had no neurological or psychiatric conditions. They also completed the Depression Anxiety Stress Scale (DASS 21) to assess well-being (Henry & Crawford, 2005). No significant differences were observed between patients and controls (depression (*df*= *18, t*=1.934, *P*=0.069) anxiety (*df*= *18, t*=1.108, *P*=0.282), and Stress (*df*= *18, t*=1.106, *P*=0.283).

All analyses of the Doors and People data were conducted in IBM SPSS Version 23. Group analyses compared neuropsychological patient and age-corrected control data raw scores using Independent *t*-tests for normally distributed data. If the Shapiro-Wilk test of normality was violated (*P*<0.05) then Mann-Whitney U Test was used for group comparison. Normality was violated for immediate verbal recall, delayed verbal recall, immediate visual recall, delayed visual recall, verbal recognition Set A, visual recognition Set A, recall and recognition discrepancy and Trail Making.
